# Characterization of OA development between sexes in the rat medial meniscal transection model

**DOI:** 10.1101/769521

**Authors:** Krishna A. Pucha, Jay M. McKinney, Julia M. Fuller, Nick J. Willett

## Abstract

**Objective:** Osteoarthritis (OA) is a chronic degenerative disease of the joints characterized by articular cartilage degradation. While there are clear sex differences in OA development in humans, most pre-clinical research has been conducted solely in male animals thus limiting the ability of these findings to be generalized to both sexes in the context of this disease. The objective of this study was to determine if sex impacts the progression and severity of OA in the rat medial meniscal tear (MMT) preclinical animal model used to surgically induce OA. It was hypothesized that differences would be observed between males and females following MMT surgery.

**Design:** A MMT model was employed in male and female Lewis rats to induce OA. Animals were euthanized 3 weeks post-surgery and EPIC-μCT was used to quantitatively evaluate articular cartilage structure and composition, osteophyte volumes and subchondral bone structure.

**Results:** Quantitative analysis of the medial 1/3 articular cartilage via EPIC-μCT showed increased cartilage thickness and proteoglycan loss in the MMT of both sexes, when compared to sham. Additionally, both male and female animals in the MMT group had increased subchondral bone mineral density and larger total osteophyte volumes due to MMT.

**Conclusion:** These data demonstrate that OA can be induced in both sexes using the rat MMT model. Moving forward, adding sex as a factor in preclinical OA studies should be standard practice in pre-clinical studies in order to elucidate more inclusive and translatable results into the clinic.

## Introduction

Osteoarthritis (OA) is the most prevalent chronic condition of synovial joints in humans and remains the leading cause of disability around the world ^1^. OA is commonly characterized by degradation of the articular cartilage in the joints with accompanying disease phenotypes of subchondral bone sclerosis, marginal tissue growths on the joint periphery, in the form of osteophytes, and synovial inflammation ^2^. In the United States alone, approximately 31 million Americans are affected by OA, and this number is only expected to increase in the next decade with the aging population ^3,4^. Numerous risk factors have been associated with disease development including age, genetics, obesity, metabolic syndrome, and sex ^5–7^.The Worldwide Health Organization (WHO) estimates that OA affects 9.6% of men and 18.0% of women over the age of 60 ^8^. Women commonly develop more severe disease phenotypes associated with higher pain levels and more substantial reduction in function and quality of life, when compared to men ^9,10^. There are numerous sex differences which can potentially be attributed to this disparity in disease incidence, but this area remains understudied and further investigation is critical in order to fully understand the mechanisms behind this phenomenon.

Two of the most well-established differences that persists in OA between sexes are body size and hormone expression. Morphological differences between males and females in articular cartilage may serve as a major contributor to differences seen in OA progression. In humans, men have on average 46.6% more cartilage volume, 32.9% more cartilage surface area and 13.3% thicker cartilage on the medial tibial plateau, relative to females ^11^. Furthermore, men demonstrate higher bone mineral density (globally) which may contribute to differences o in subchondral bone mineral density loss associated with OA progression ^11,12^. Consideration of whole joint biomechanical parameters, particularly in the knee, may also play a role in this disparity of disease incidence as women demonstrate increased mechanical loading on the knee due to differences in lower limb alignment ^13^. Other sex differences in animal models also exist, such as weight and growth rates, which could impact the development of OA, as they do in humans ^14,15^. While these anatomical and structural differences potentially have a major impact on differences in OA progression between sexes there are numerous other factors which can also influential.

Men and women demonstrate markedly different expression of hormones and tend to have different physiological responses to the signaling cues generated by these hormones. Numerous reports have highlighted the varied mechanisms of action and receptor activity that persists for testosterone and estrogen between males and females, but notable differences have also been described for other hormones, including leptin ^16–19^. The role that estrogen plays in OA has been well defined, as decreased expression of this hormone directly correlates to enhanced OA incidence ^20–22^. This is demonstrated in the phenomenon commonly referred to as ‘menopausal arthritis’, describing the dramatic increase in the prevalence and severity of OA in women around the age of 50, coinciding with the onset of menopause in this same population ^23^. Furthermore, hormone replacement therapy (HRT) has shown promise in slowing OA progression in animal models of OA as well as in the clinical setting ^20–22^. There is substantial data documenting the differences of OA progression between sexes in humans which have been extensively reviewed previously ^24-26^. However, there remains a disparity of knowledge in the way sex effects OA progression in preclinical animal models. These models create the foundation for clinical research to build upon, are commonly used in mechanistic studies of the disease and are used to develop and test novel therapeutic approaches. Thus it is vital to understand any potential differences in disease development between the sexes in these pre-clinical models.

Animal models of OA broadly range from small animals like the mouse destabilization of the medial meniscus (DMM) and rat medial meniscal transection (MMT) to larger animal like goats, guinea pigs and even horses. This spectrum of models provides a practical pipeline to study both mechanisms of disease development and progression and to evaluate therapeutic efficacy of new treatments ^27,28^. Even though sex is a critical factor in OA development in humans, most pre-clinical studies do not account for or address sex differences when using animal models to study mechanisms of OA development or test new therapies ^24^. Additionally, many published review articles that discuss preclinical animal models of OA do not address the sex of the animals used for the respective studies or the effect sex can have on OA progression in these models; this is in large part is due to a pervasive lack of studies that have investigated sex differences in these pre-clinical models ^25–28^.

In the current study we utilized one of the most common pre-clinical animal models, the Lewis rat. This animal model has a thick cartilage surface, for a rodent, and OA can easily be induced through a medial meniscus transection (MMT) surgery, making it a commonly used model ^29^. Though sex differences in OA animal models have been addressed to a small extent, such as with the mouse DMM model and cynomolgus monkey, there is limited understanding of the potential presence of any sex difference in OA development in rat models employed for OA research even though these models are utilized in a fourth of all published reports using animal models to study OA ^14,27,30^. These models, typically done in a single sex, do not distinguish between sexes or therapeutic efficacy between sexes. Accounting for sex differences in these disease models is critical as they can impact the clinical aspect of OA research. Furthermore, recent NIH mandates also require that sex be considered when using animal models to study disease in order to enable researchers to draw conclusions that can be generalizable to all patients ^31^. The objective of the current study was to characterize the presence of sex differences, if any, in OA development for the MMT rat model. It was hypothesized that a difference would be seen in the progression of OA between male and female Lewis rats.

## Methods

### Surgical Methods

All animal care and experiments were conducted in accordance with the institutional guidelines of the Atlanta Veteran Affairs Medical Center (VAMC) and experimental procedures were approved by the Atlanta VAMC Institutional Animal Care and Use Committee (IACUC) (Protocol: V002-18).

Male (n=16) Lewis rats, weighing 300-350 g, and female (n=15) Lewis rats, weighing 275-300 g (strain code: 004; Charles River), were ordered and acclimated to the environment for one-week post arrival to the VAMC. The MMT pre-clinical model, a model of surgical instability, was conducted to induce OA in rats as previously described ^25^. Briefly, isoflurane inhalation was used to anesthetize the animals and SR buprenorphine (ZooPharm, Windsor, CO) was injected subcutaneously at 1mL/kg as an analgesic during the procedure. Skin was shaved and sterilized at the site of surgery. An initial incision was made in the skin along the medial aspect of the femora-tibial joint. A blunt dissection was then made in this area to expose the medial collateral ligament (MCL) and the MCL was transected to expose the meniscus. A full thickness cut of the meniscus was made at the narrowest point to complete the MMT procedure. The wound, in the musculature, soft tissue and skin, was closed following surgery using 4.0 vicryl sutures and staple clips. Sham surgeries were also preformed, where the MCL was transected but not the meniscus.

Post-surgery, the rats were injected subcutaneously with 10 mL of Lactated Ringer’s solution and a liquid bandage and Metronidazole paste were applied to the incision area to aid recovery and deter the animals from further irritating the wound. At 7 days post-surgery, staples were removed from the incision area.

The rats were euthanized via CO2 asphyxiation 3 weeks post-surgery and the hind limbs were collected. At this time point, it has been established that early stage OA has developed in the animals ^25^. The samples were stored in 10% neutral buffered formalin for fixation.

### Sample Preparation

To prepare the samples for analysis, all muscle and connective tissue surrounding the joint, along with the synovium, was removed from the limbs and the femur was disconnected from the tibia to expose the cartilaginous surface of the tibial condyle. The tibiae were removed from formalin and stored in PBS until scanning. Immediately before scanning, the tibiae were immersed in 30% Hexabrix 320 contrast reagent (NDC 67684-5505-5, Guerbet, Villepinte, France) for 30 min at 37 °C ^32^.

### EPIC-μCT analysis of articular cartilage, osteophytes and subchondral bone

To quantitatively analyze the structure and composition of the articular cartilage and the morphology of osteophytes and the subchondral bone, equilibrium partitioning of an ionic contrast agent based micro-computed tomography (EPIC-μCT) analysis was utilized. The proximal tibia was scanned with a Scanco µCT 40 (Scanco Medical, Brüttisellen, Switzerland). Scan parameters were: 45 kVp, 177 μA, 200 ms integration time, 16 μm voxel size, ∼ 27 min scan time ^32^. Raw scan data were automatically reconstructed into two-dimensional (2D) grayscale tomograms and orthogonally transposed to yield coronal sections.

Coronal sections for the articular cartilage were evaluated across the full length of the tibial condyle and analyzed specifically in the medial 1/3 region of the medial tibial condyle as this region has been shown to demonstrate the most damage in the MMT-induced OA model ^33^. Contours were constructed to isolate the articular cartilage surface and threshold values of 110 and 435 mg hydroxyapatite per cubic centimeter (mg HA/cm^3^) were used to separate cartilage from bone and air. The Scanco µCT 40 generated values for articular cartilage volume, thickness and attenuation. Cartilage attenuation is inversely related to sulfated glycosaminoglycan (sGAG) content ^32^. Cartilage degradation leads to lower levels of sGAG in the cartilage leading to higher concentrations of Hexabrix 320 contrast reagent in the extra cellular matrix of the cartilage. Higher Hexabrix corresponds with lower sGAG and higher attenuation levels ^32^.

Osteophytes are marginal tissue growths that form on the medial aspects of the tibial condyle and are a common symptom of OA ^34^. Osteophyte volumes were evaluated using coronal sections generated by the scanning of the tibiae ^35,36^. Cartilaginous osteophytes were isolated from surrounding air and bone using threshold values of 110-435 mg HA/cm^3^. Mineralized osteophytes were isolated from the surrounding cartilage and soft tissue using threshold values of 435-1000 mg HA/cm^3^.

Coronal sections for the subchondral bone were evaluated across the full length of the tibial condyle and the medial 1/3 was isolated for analysis. Threshold values of 435-1000 mg HA/cm^3^ were used to isolate the bone from the surrounding tissue. The Scanco µCT 40 generated values for subchondral bone volume, thickness, attenuation and porosity.

### Surface roughness analysis of articular cartilage

To quantify surface roughness, coronal sections generated by scanning the tibiae were exported as 2D TIFF images and imported into MATLAB R2016a (MathWorks, Natick, MA). A custom code created a 3D surface with these images by scanning section images sequentially and fitting the 3D surface along a previously generated 3D polynomial surface: fourth order along the ventral-dorsal axis and second order along the medial-lateral axis ^35^. The surface roughness was quantified as the root mean square of differences between the 3D surface created with the exported TIFF images and the polynomial fitted surface. Surface roughness was calculated across the medial 1/3 region of the tibial plateau in accordance where the predominant damage is found in the MMT model ^38^. Threshold values of 110 and 435 (mg HA/cm^3^) were used to separate cartilage from air and the underlying subchondral bone.

### Histology

Tibiae were decalcified with formic/citrate decalcifying solution (Newcomer Supply 10429C, Middleton, WI) for 7 days. They were then dehydrated and processed with paraffin-imbedded blocks. The samples were then sectioned into 5 μm-thick slices and stained with hematoxylin and eosin (H&E; Fisherbrand™ 517-28-2, Waltham, MA) or safranin O and fast green (Saf-O; Electron Microscopy Sciences® 20800, Hatfield, PA), following manufacturer protocols.

### Statistical analysis

All figures are presented as the calculated mean +/- standard deviation (SD). Significance was determined using two-way analysis of variance (ANOVA) test between groups with post-hoc Tukey Honest analysis for articular cartilage and subchondral bone. Significance for osteophyte data was determined using two-way ANOVA with a nonparametric post-hoc Bonferroni analysis, due to the non-uniform distribution of osteophyte volumes. For all statistical analyses, significance was set at *p* < 0.05. Data were analyzed and significance determined using GraphPad Prism software version 6.0 (GraphPad Software Inc., La Jolla, CA).

## Results

### Qualitative analysis of MMT induced OA

Histological staining of tibial sections by H&E and Saf-O was used to qualitatively assess the extent of damage caused by the MMT model. Samples from the sham groups for both males and females depicted intact cartilage with smooth superficial surfaces and relatively uniform proteoglycan content across the medial tibial condyle. Osteophytes were not apparent on the marginal edges of the joint for either of the sham groups. MMT samples from both sexes depicted large amounts of proteoglycan loss, apparent in qualitatively less Saf-O (red) staining, especially in the medial 1/3 region of the medial tibial condyle. Both Saf-O and H&E staining showed fibrillations and lesions on the cartilage surface in both males and females with MMT. MMT samples in both sexes also showed the presence of osteophytes along the medial edge of the tibiae (Fig. 2). EPIC-μCT images depicted similar qualitative observations in cartilage composition and structure and osteophyte formation as seen in the histological analysis (Fig. 2).

**Figure 1:**
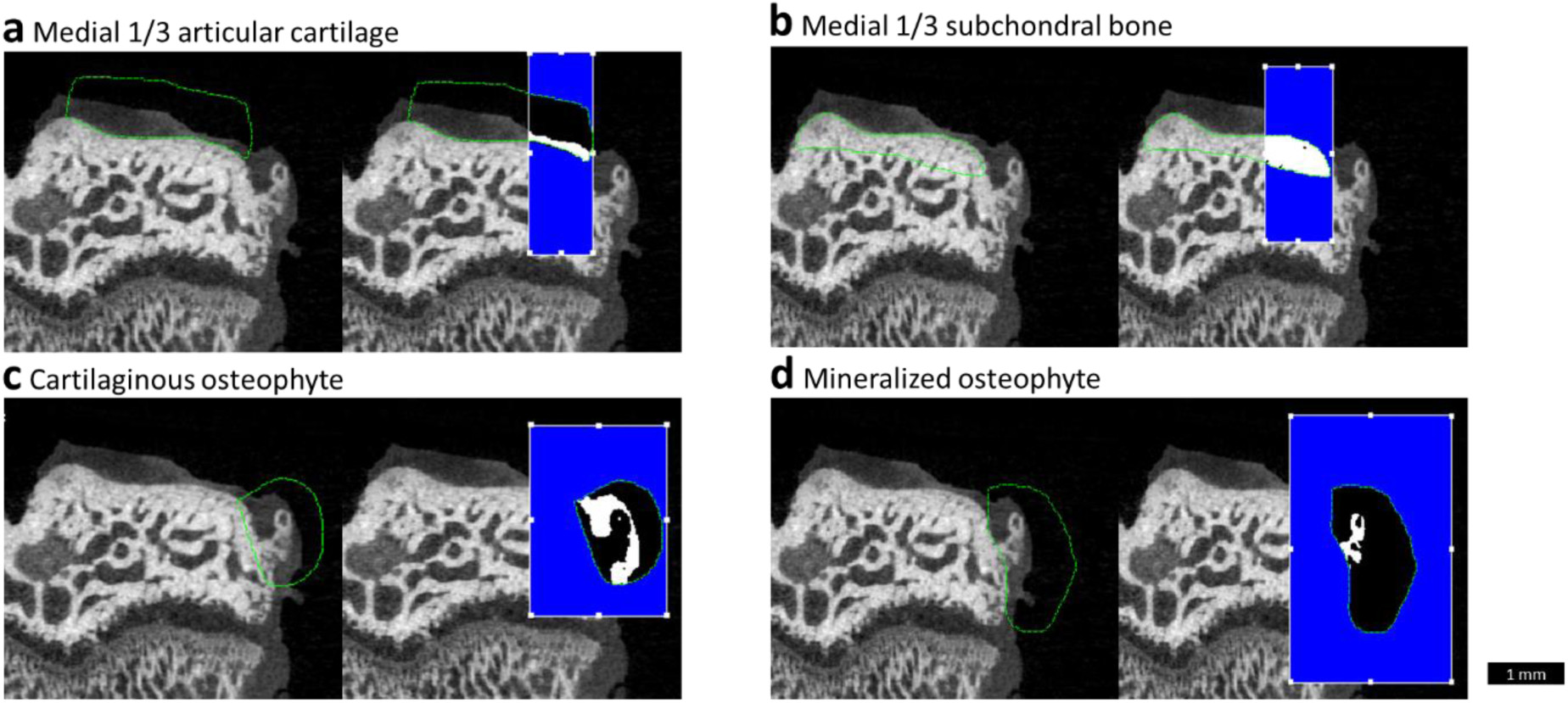
Representative EPIC-μCT images depicting contoured volume of interest (VOI) for medial 1/3 articular cartilage and subchondral bone and mineralized and cartilaginous osteophytes. Representative coronal sections showing a rat medial tibial condyle with contours outlined in green and the medial 1/3 VOI for each respective aspect of the tibia highlighted in white for total medial **(a)** articular cartilage and **(b)** subchondral bone. Representative coronal sections with contours outlined in green and analyzed volumes highlighted in white for **(c)** cartilaginous and **(d)** mineralized osteophyte in MMT joint. Scale bar is universal for all representative images.

**Figure 2:**
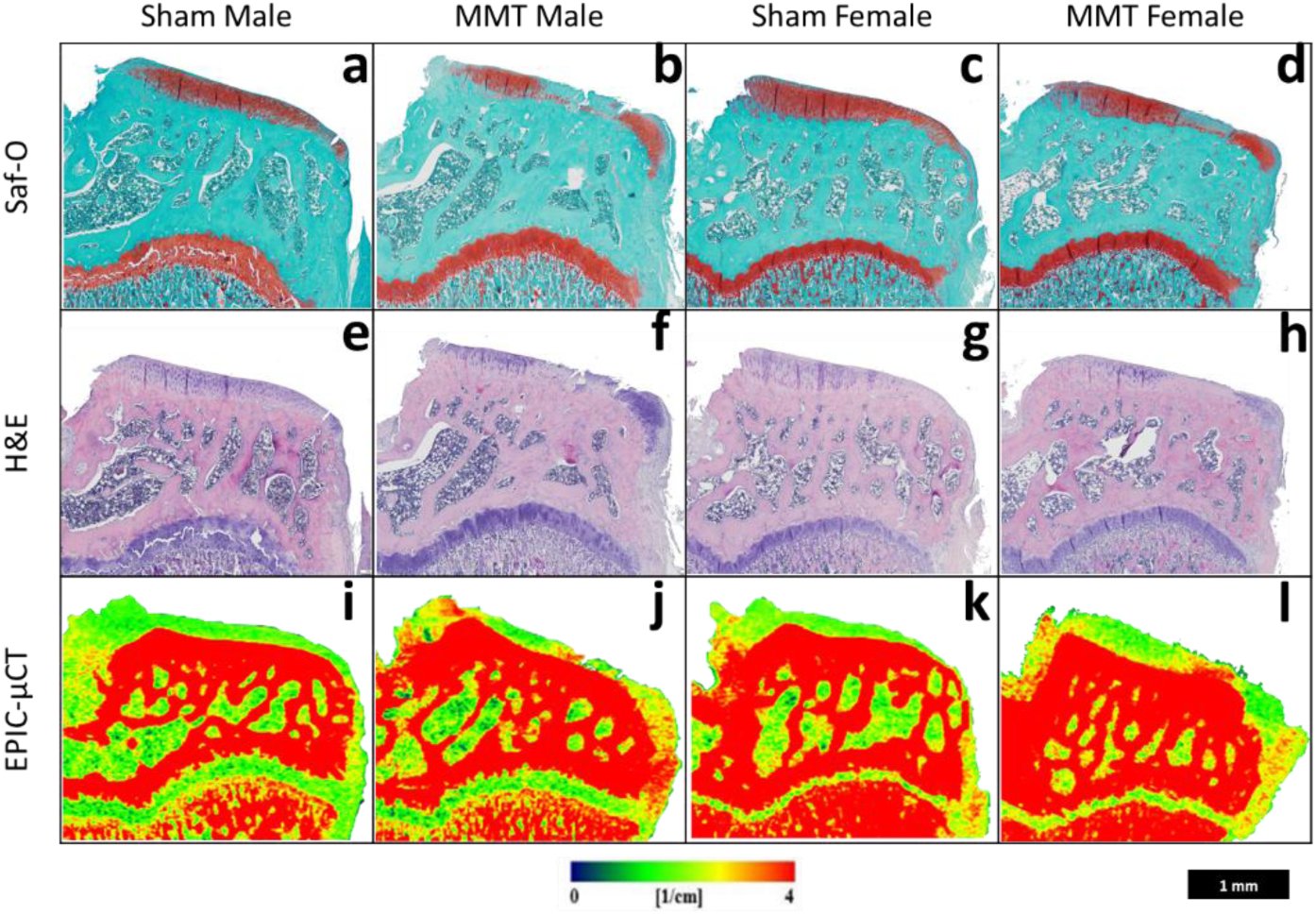
Histological and EPIC-μCT representative coronal sections of rat medial tibial condyle for sham and MMT joints for both male and female animals. **(a-d)** Saf-O and **(e-h)** H&E stained histological sections from a rat medial tibia showing the presence of fibrillations on the articular cartilage surface, osteophyte formation, proteoglycan loss (as depicted by absence of red coloration in **(a-d)** Saf-O stained samples) and loss in cell viability in articular cartilage layer (as depicted by absence of hematoxylin (violet coloration) in **(e-h)** H&E stained samples) for MMT **(b, f)** male and **(d, h)** female animals. Sham **(a, e)** male and **(c, g)** females animals had no damage, cartilage degeneration, or osteophyte formation. EPIC-μCT representative images show the presence of osteophytes, fibrillations and cartilage degradation (attenuation is inversely proportional to proteoglycan content) in **(j, I)** MMT samples and no damage in **(i, k)** sham samples. All images are oriented with the medial aspect of the tibia on the right. Scale bar (bottom right corner) is universal for images.

### EPIC-μCT quantitative analysis of medial 1/3 articular cartilage

To quantitatively analyze OA progression in male and female rats in the MMT model, the medial 1/3 region of the articular cartilage of the medial tibial condyle was evaluated using EPIC-μCT. Cartilage damage has been shown to initiate in this region of the tibial condyle and is detectable by EPIC-μCT at the 3 week post-surgery time point in the MMT model^33^. Attenuation values for males and females both yielded increased values in the MMT groups compared to their respective sham groups, indicative of a decrease in proteoglycan content in the MMT condition relative to the sham (Fig. 3a). When comparing sham and MMT groups, both male and female animals showed significant increases in cartilage volume and thickness in the MMT group, relative to the respective sham groups (Fig. 3b&c). Furthermore, in the male animals, both in the sham and MMT groups, the cartilage volume and thickness parameters were increased relative to the female animals, respectively (Fig. 3b&c). To account for the inherent size differences between male and female animals, cartilage volume and thickness values were weight normalized to permit direct comparison of the magnitude changes between sham and MMT animals between the two sexes (Supp. Fig. 1). Upon normalization of volume and thickness values by the mass of each animal both males and females had significant increases in medial 1/3 cartilage volume and thickness, when comparing shams and MMT for each sex independently. Comparing between sexes, the MMT females had a significantly increase in the normalized cartilage thickness compared to MMT males.

**Figure 3:**
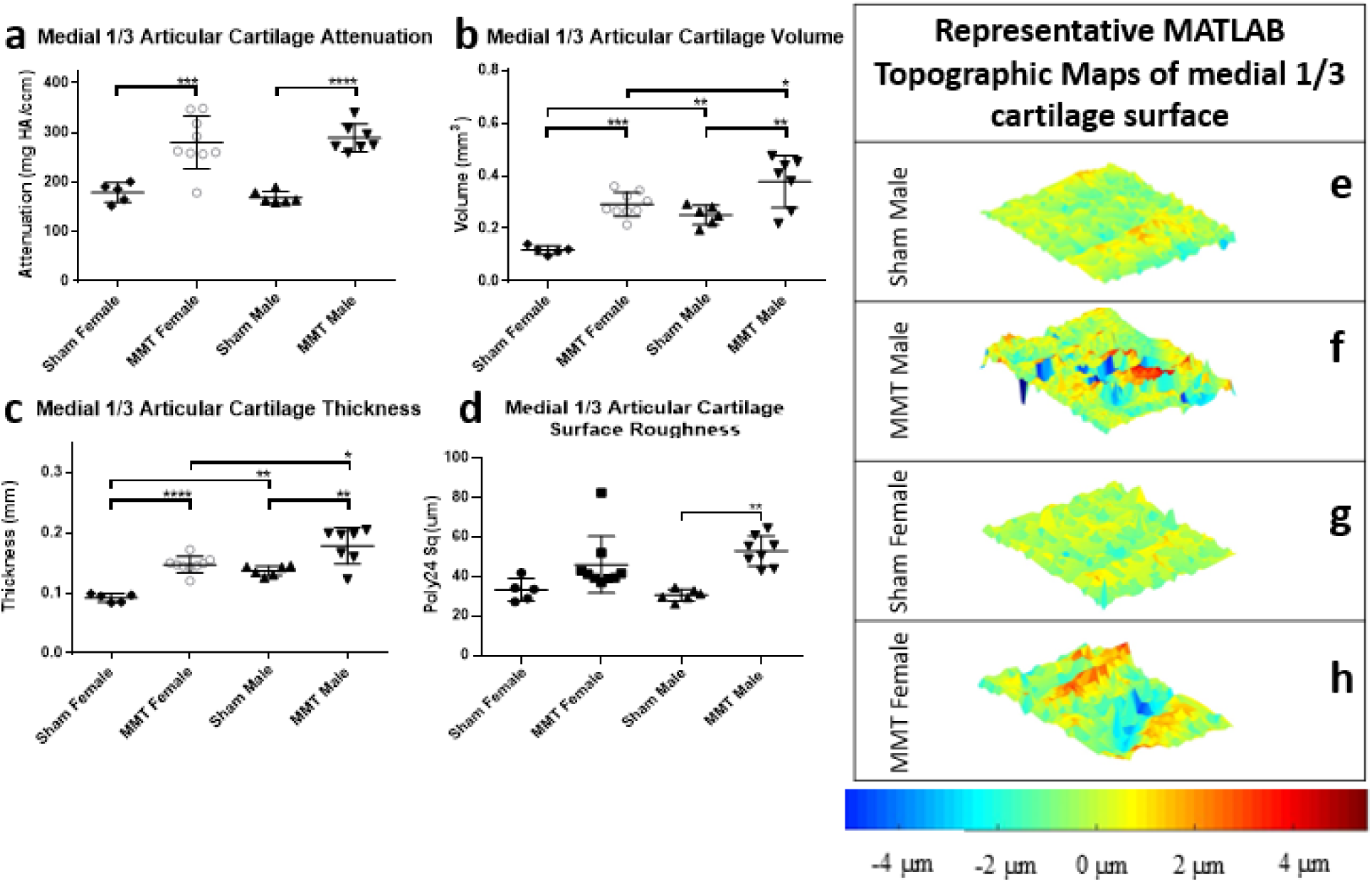
EPIC-μCT quantitative analysis of medial 1/3 articular cartilage in sham and MMT tibiae from both sexes. **(a)** Medial 1/3 articular cartilage attenuation increased in MMT joints compared to sham tibiae for both male and female animals with no significant differences between male and female animals. Medial 1/3 articular cartilage **(b)** volume and **(c)** thickness increased in MMT tibiae compared to sham joints for both male and female animals; males had higher values than females when comparing MMT males to MMT females and sham males to sham females for both parameters. Medial 1/3 articular cartilage surface roughness significantly increased in the MMT male animals compared to male sham animals. **(e-h)** MATLAB generated representative topographic maps of medial 1/3 cartilage surface depict the deviation of a corresponding sample’s cartilage surface renderings from a standardized 3D polynomial surface. Topographic maps show qualitatively increased surface roughness in MMT **(f)** male and **(h)** female animals compared to **(e, g)** shams. Data shown as mean +/- SD. *n* = 5 for sham females, *n* = 9 for MMT females, *n* = 6 for sham males and *n* = 7 for MMT males. *p < 0.05; **p ≤ 0.01; ***p < 0.001; ****p ≤ 0.0001.

MATLAB analysis of cartilage surface roughness showed increased roughness values for male animals but not female animals in the MMT group compared to the sham group (Fig. 3d). Qualitative analysis of the cartilage surface roughness, in the form of topographic maps, for all study groups were completed by subtracting individual 3D polynomial surfaces from the corresponding cartilage surface renderings yielded (Fig. 3 e-h).

### EPIC-μCT quantitative analysis of osteophytes

EPIC-μCT was used to quantitatively asses the volumes of osteophytes, defined as thickenings and partial mineralizations of the tissue at the marginal edge of the medial tibial plateau ^34^. Both sexes yielded significantly larger cartilaginous osteophyte volumes in their respective MMT groups compared to shams. Furthermore, male animals had significantly larger cartilaginous osteophytes in the MMT condition compared to female MMT animals (Fig 4a). For the mineralized osteophyte volumes, a similar result was observed, in that the MMT males exhibited significantly increased volumes relative to the MMT females (Fig. 4b). The MMT male group also had significantly higher mineralized osteophyte volumes, relative to the sham; whereas no significant difference was observed in the female MMT group. Total osteophyte volumes, which were calculated as a sum of cartilaginous and mineralized osteophyte volumes, showed significantly increased values in the MMT group for both male and female groups, relative to the shams. Furthermore, and consistent with the increased cartilaginous osteophyte volumes, male MMT total osteophytes volumes were significantly higher than that of MMT females (Fig 4c). To again account for the inherent size differences between male and female animals all osteophyte data sets were normalized between the sexes by dividing the respective volume measurements by the mass of the animals (Supp. Fig. 2). This normalized analysis showed significantly increased values for all osteophyte parameters (cartilaginous, mineralized and total) in the male and female MMT groups compared to the respective shams. In addition, the normalized data showed significantly greater cartilaginous and total osteophyte volumes in male MMT animals compared to female MMT animals.

**Figure 4:**
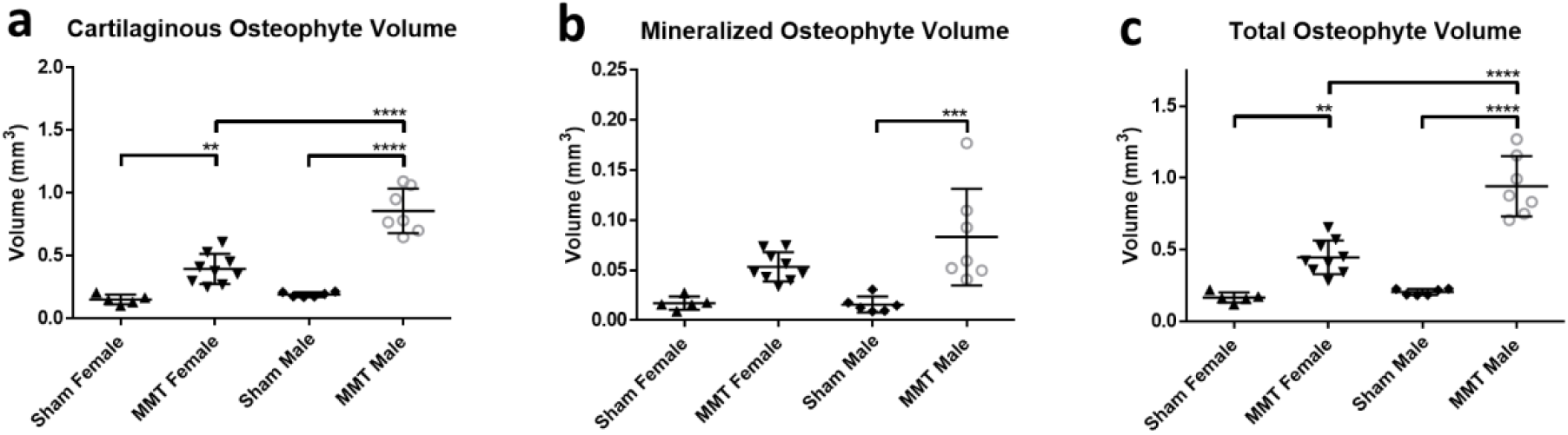
EPIC-μCT quantitative analysis of osteophyte formation on the medial tibiae of sham and MMT joints from both sexes. **(a)** Cartilaginous and **(c)** total osteophyte volumes were significantly increased in MMT joints compared to shams for both sexes with MMT male volumes significantly higher than MMT females. **(b)** Mineralized osteophyte volumes significantly increased in MMT male animals compared to sham males. Data shown as mean +/- SD. *n* = 5 for sham females, *n* = 9 for MMT females, *n* = 6 for sham males and *n* = 7 for MMT males. *p < 0.05; **p < 0.01; ***p < 0.001; ****p < 0.0001.

### EPIC-μCT quantitative analysis of subchondral bone

EPIC-μCT was used to quantify the effects of MMT surgery on changes to the subchondral bone in the medial 1/3 region of the subchondral bone. Analysis of subchondral bone volume in females showed increased thickness values in the MMT group relative to sham group, while no differences were detected in male animals (Fig. 5a). Furthermore, for the thickness of the subchondral bone there was no detectable differences between sham and MMT for either male or female animals (Fig. 5b). Subchondral bone mineral density was significantly increased in the MMT groups relative to the sham group in both males and females (Fig. 5c). To account for the inherent size differences between male and female animals, as was done with articular cartilage and osteophyte analyses, subchondral bone volume and thickness values were normalized by mass to permit direct comparison of the relative magnitude changes between sham and MMT animals between the sexes (Supp. Fig. 3). Normalized data showed an increase in medial 1/3 subchondral bone volume and thickness only in female MMT animals whereas the male MMT groups and male sham groups were statistically identical.

**Figure 5:**
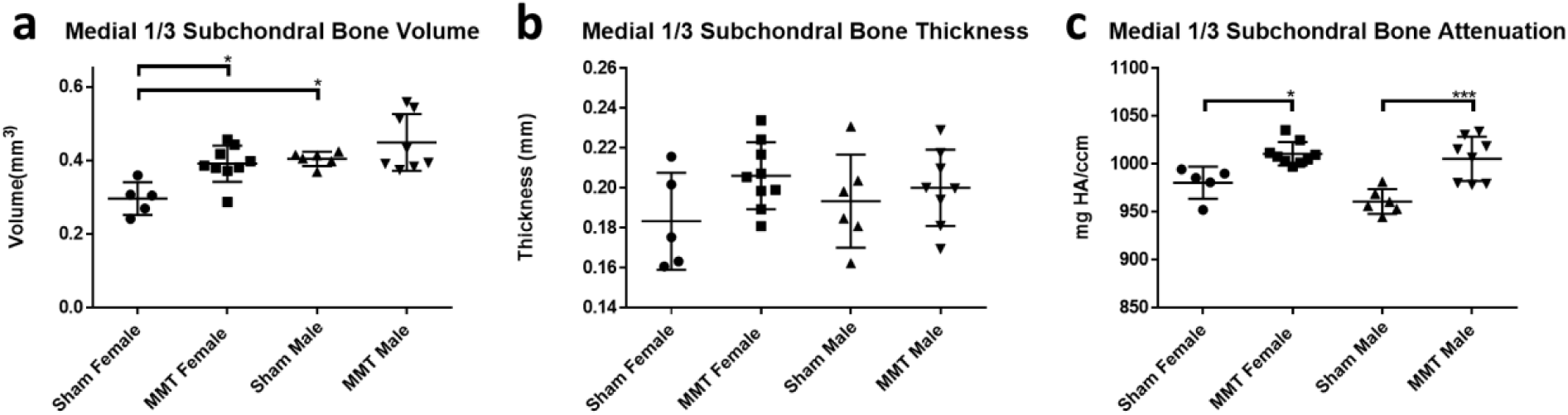
EPIC-μCT quantitative analysis of the medial 1/3 subchondral bone in sham and MMT tibiae from both sexes. **(a)** Medial 1/3 subchondral bone volumes significantly increased in MMT female animals compared to sham female animals. Sham male animals had significantly higher volumes than sham female animals. **(b)** Medial 1/3 subchondral bone thickness had no significant changes in MMT animals compared to sham for both sexes and no significant differences were detected between male and female animals. **(c)** Medial 1/3 subchondral bone attenuation (density) increased in MMT joints compared to sham joints for both male and female animals with no significant differences between male and female animals. Data shown as mean +/- SD. *n* = 5 for sham females, *n* = 9 for MMT females, *n* = 6 for sham males and *n* = 7 for MMT males. *p < 0.05; **p < 0.01; ***p ≤ 0.001; ****p ≤ 0.0001.

## Discussion

Pre-clinical animal models of OA are commonly used to elucidate the underlying mechanisms of the disease and to assess the efficacy of therapeutics being developed to treat OA. For these pre-clinical models, it is critical to be able to characterize and consider the major variables that can affect OA progression. Accounting for how these factors change over the course of disease progression permits comprehensive conclusions to be drawn therefore allowing for more effective translation towards the clinic. Differences in OA development and progression between the two sexes in pre-clinical models is a critical factor to consider and is one that remains understudied. Sex based variables and differences may directly influence the efficacy of treatment and progression of disease and therefore must be considered to ensure successful clinical translation of therapeutics developed on the pre-clinical side. This motivates further study into OA progression in both males and females in pre-clinical animal models. The objective of the current study was to characterize the effect of sex differences in OA progression in the MMT pre-clinical Lewis rat model.

The results of the current study showed that males possessed increased cartilage volume and thickness relative to females, as was observed by the difference in both articular cartilage volume and thickness in the sham groups of males and females; these pre-clinical structural findings are consistent with previously published data from humans and other animal models ^11,37^. Animals were not weight or age matched due to the generally lower weight of female animals at a given a age; animal weight varied significantly between the sexes with males having a mean weight of 325 grams and females having a mean weight of 211 grams. This is a potential limitation of the study. Regardless of the differences in cartilage morphology and body mass, both males and females developed OA after MMT surgery, as evidenced by significant increases in cartilage attenuation (indicative of proteoglycan loss), volume and thickness in the MMT groups of both male and female animals, when compared to their respective sham groups. This is also consistent with the mouse DMM model, in which previous studies have also showed OA development in both sexes; though in the DMM model, males showed increased disease severity ^30^.

The current study utilized µCT as a means to quantitatively assess OA progression between sexes. This approach allowed for the size differences between the animals to be taken into consideration and to be used to normalize the data, a constraint that the previously mentioned studies have not been able to include due to the limitations of histomorphometric analysis. Accounting for these inherent size differences when comparing sexes in this animal model is important as the MMT is a post-traumatic OA model and much of the damage observed can be attributed to the mechanical load placed on the cartilage surface in the animals ^38^. Previous studies have suggested that male mice exhibit a higher degree of damage to the articular cartilage layer ^30,39^. However, in the study we presented here, we observed similar increases in magnitudes of normalized articular cartilage volume in both male and female MMT cartilage volumes compared to the corresponding shams.

The formation of osteophytes in the medial aspect of the tibia is also an indicator of OA development and a key feature that is replicated in the pre-clinical rat MMT model. Though current understanding of osteophyte formation is limited, generally, osteophytes are defined as marginal tissue growths that form on the medial aspects of the tibial condyle and consist of cartilaginous and mineralized parts ^40^. Increased body mass, which leads to increased loads on the knees, has been shown to increase osteophyte formation in the knee joint ^41,42^. Quantitative analysis in the current study showed a significant increase in total osteophyte volume in both male and female MMT animals; male animals showed larger total osteophyte growth compared to females while MMT females did not show any significant increase in mineralized osteophyte volumes compared to shams. In previous studies done with the mouse DMM model, increased osteophyte growth was observed in male animals compared to females, but the effect of the confounding variable of weight and associated mechanical load changes on osteophyte formation was not addressed ^43^. Additionally, when characterizing osteophyte progression between sexes in the DMM model, there was no discrimination between key compositional components of osteophytes, including the mineralized and cartilaginous portions ^43^. However, when weight was used to normalize for osteophyte volumes in the current study, females did demonstrate significant increases in mineralized formations. However, male animals still yielded significantly larger total osteophyte volumes than female animals. The differences in osteophyte development may be attributed to the difference in weight between the two sexes, consistent with the general understanding of osteophyte formation. It is also possible that the increased joint size and size of the tibial condyle in male animals may provide additional room for osteophytes to form^44^. While these differences are important to consider, both sexes developed osteophytes with mineralized and cartilaginous portions underscoring the utility of this model in both males and females and the need to utilize both sexes when studying these features. Mineralized osteophyte formations are of particular importance in the clinic as X-rays are often unable to differentiate between cartilage and soft tissue since they are similar in composition and look identical when examined via X-ray ^45^.

The subchondral bone is another critical tissue and component of joint health that can be impacted during OA development and progression. As the disease progresses the subchondral bone will often undergo both hardening and thickening (sclerosis) ^46^. Previous studies that have characterized the effects of sex on OA progression in the mouse DMM model focused primarily on changes to the cartilage and not the subchondral bone ^30,39^. In our current study, we assessed subchondral bone changes by analyzing subchondral bone morphometry and composition using μCT. μCT data showed that both male and female animals had an increase in bone mineral density after MMT surgery, suggesting the induction of sclerosis in this underlying bone for these animals. However, no differences were noted for either volume or thickness parameters, even when accounting for weight. While increased bone mineral density was observed in the current study, previous studies involving the male MMT models have demonstrated that significant morphological changes in the subchondral bone tend to emerge at later time points ^47^. The MMT model used for OA induction yielded expected subchondral bone morphologies, and both males and females developed these changes in the subchondral bone. Later timepoints potentially would show more pronounced changes in the subchondral bone.

This study demonstrated that OA in the MMT model can be induced in both sexes and that overall, the development of OA in both sexes of this preclinical model was largely similar. Analyzing individual aspects of joint health allowed for comprehensive quantitative analysis of disease development in three dimensions and showed changes in articular cartilage, osteophytes and subchondral bone. The analyses conducted showed factors associated with OA development which are compositional in nature and independent of weight, such as attenuation and bone density, did not differ between sexes; however, structural parameters associated with OA development, such as mineralized osteophyte and medial 1/3 subchondral bone and articular cartilage volumes which potentially arise from differences in weight, should be given extra consideration when utilizing this model with both sexes. Additionally, hormonal and certain morphological differences could also be included in assessment of disease progression as these factors could also potentially contribute to the results observed in this study. Using both sexes in the MMT model, will allow for studies to account for differences in disease development that might arise due to these factors, in addition to weight. As this preclinical model is often used to test the efficacy of therapeutics, this study demonstrates that it is appropriate to use both sexes in pre-clinical studies to ensure that therapies can slow the progression of OA in both sexes.

## Supplemental Figures

**Supplemental Figure 1:**
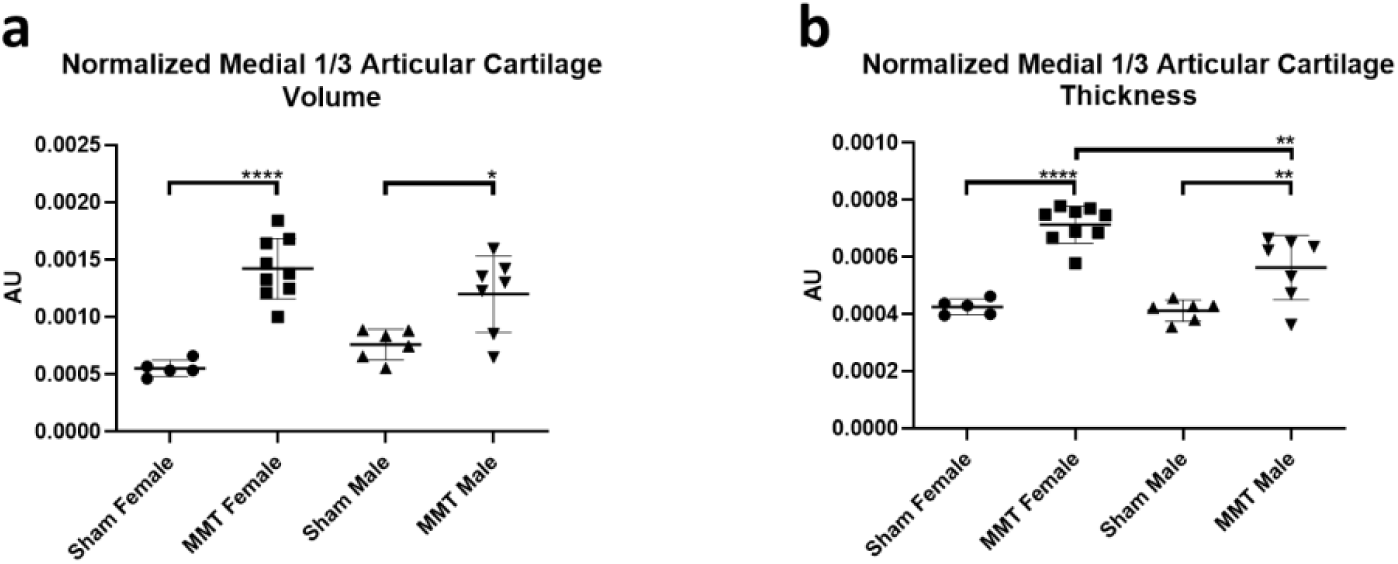
Mass normalized EPIC-μCT quantitative analysis of osteophyte formation on medial tibia of sham and MMT joints from both sexes. Mass normalized values were obtained by dividing the value from each sample by the sample’s corresponding mass measured on the day of surgery. **(a)** Mass normalized medial 1/3 articular cartilage volumes increased in MMT joints compared to sham joints for both male and female animals with no significant differences between male and female animals. **(b)** Mass normalized medial 1/3 articular cartilage thickness increased in MMT joints compared to sham joints for both male and female animals with MMT female animals showing significantly greater volume compared to MMT males. Data shown as mean +/- SD. *n* = 5 for sham females, *n* = 9 for MMT females, *n* = 6 for sham males and *n* = 7 for MMT males. *p < 0.05; **p ≤ 0.01; ***p ≤ 0.001; ****p ≤ 0.0001.

**Supplemental Figure 2:**
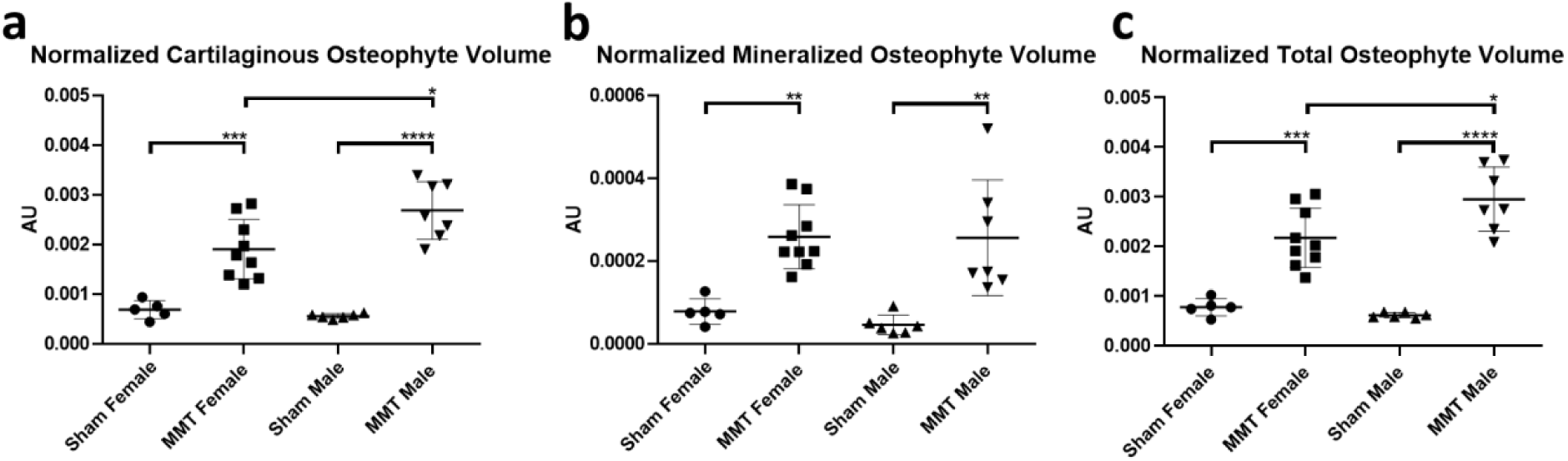
Mass normalized EPIC-μCT quantitative analysis of osteophyte formation on medial tibia of sham and MMT joints from both sexes. Mass normalized values were obtained by dividing the value from each sample by the sample’s corresponding mass measured on the day of surgery. Mass normalized **(a)** mineralized and **(c)** total osteophyte volumes increased in MMT joints compared to sham joints for both male and female animals with MMT male animals showing significantly greater volume compared to MMT females. **(b)** Mass normalized mineralized osteophyte volumes increased in MMT joints compared to sham joints for both male and female animals with no significant differences between male and female animals. Data shown as mean +/- SD. *n* = 5 for sham females, *n* = 9 for MMT females, *n* = 6 for sham males and *n* = 7 for MMT males. *p < 0.05; **p ≤ 0.01; ***p ≤ 0.001; ****p ≤ 0.0001.

**Figure 5:**
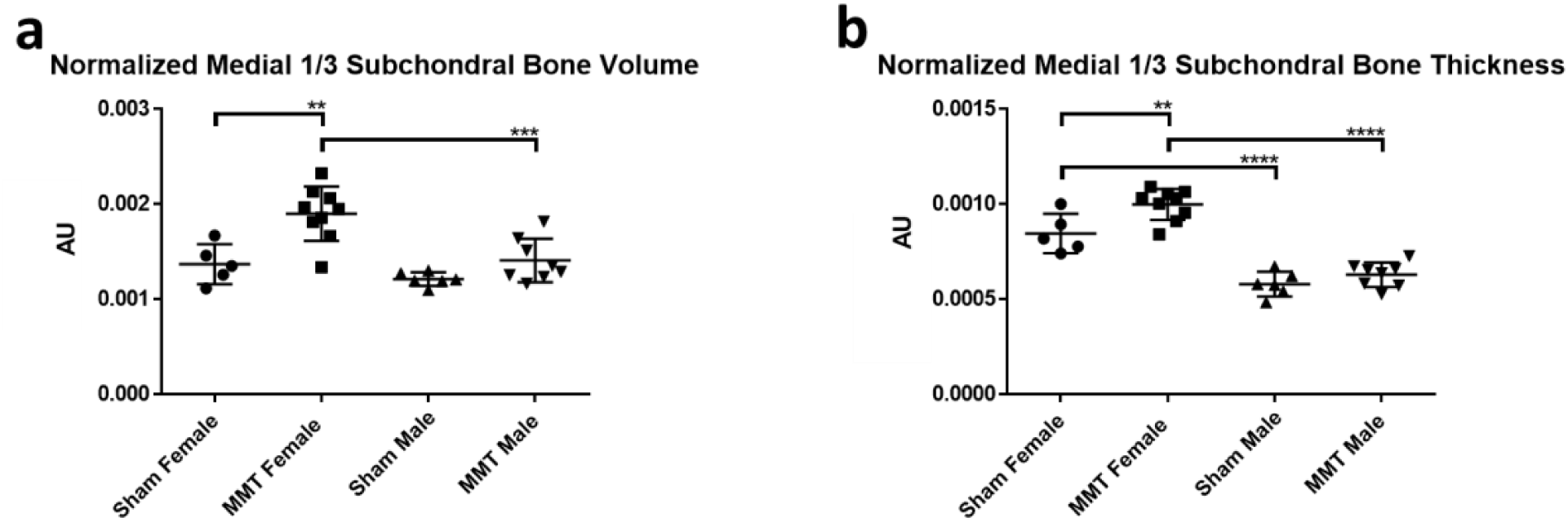
Mass normalized EPIC-μCT quantitative analysis of medial 1/3 subchondral bone in sham and MMT joints from both sexes. **(a)** Mass normalized medial 1/3 subchondral bone volumes significantly increased in only MMT female animals compared to sham female animals. **(b)** Mass normalized medial 1/3 subchondral bone thickness significantly increased in only MMT female animals compared to sham female animals with female with females showing larger values than males when comparing MMT males to MMT females and sham males to sham females. Data shown as mean +/- SD. *n* = 5 for sham females, *n* = 9 for MMT females, *n* = 6 for sham males and *n* = 7 for MMT males. *p < 0.05; **p ≤ 0.01; ***p≤ 0.001; ****p ≤ 0.0001.

